# Independent synaptic inputs to motor neurons driving antagonist muscles

**DOI:** 10.1101/2022.08.18.504426

**Authors:** Daniele Borzelli, T.M.M. Vieira, A. Botter, M. Gazzoni, F. Lacquaniti, A. d’Avella

## Abstract

The CNS may produce the same endpoint trajectory or torque profile with different muscle activation patterns. What differentiates these patterns is the presence of co-contraction, which does not contribute to joint torque generation but allows to modulate mechanical impedance. Whether co-contraction is controlled through the same synaptic input to motor neurons involved in generating joint torque is still unclear. We hypothesized that co-contraction is controlled through a specific synaptic input, independent from that underlying the control of torque. To test this hypothesis, we asked participants to concurrently generate multi-directional isometric forces at the hand and to modulate the co-contraction of arm muscles to displace and stabilize a virtual end-effector. The firings of motor units were identified through decomposition of High-Density EMGs collected from two antagonist muscles, Biceps Brachii and Triceps Brachii. We found significant peaks in the coherence between the neural drive to the two muscles, suggesting the existence of a common input modulating co-contraction across different exerted forces. Moreover, the within-muscle coherence computed after removing the component synchronized with the drive to the antagonist muscle or with the exerted force revealed two subsets of motor neurons that were selectively recruited to generate joint torque or modulate co-contraction. This study is the first to directly investigate the extent of shared versus independent control of antagonist muscles at the motor neuron level in a task involving concurrent force generation and modulation of co-contraction.

**Significance Statement:** How the CNS coordinates the activity of antagonist muscles to modulate limb mechanical impedance is still unclear. We hypothesized that a common synaptic input, shared by the motor neurons pools of antagonist muscles, and independent from the inputs underlying force generation, regulates co-contraction. We then analyzed the coherence between the firing trains of motor neurons to assess whether a common input drives antagonist muscles only during tasks requiring co-activation for impedance but not for force generation. Results highlighted the existence of separate neural pathways underlying the control of joint torque or impedance. Scientifically, this study addressed an important gap in understanding how neural drive is delivered to antagonist muscles, disentangling the control of muscles for joint torque or impedance modulation.

## Introduction

The musculoskeletal system, owing to its redundancy (Bernstein, 1967), may produce the same joint trajectory or torque profile with a multitude of combinations of muscle activation patterns differing in the amount of co-contraction, i.e. the co-activation of multiple muscles generating joint torques with opposite signs and thus no endpoint force. Co-contraction helps modulating impedance (Hogan, 1984; Milner, 2002) and therefore reducing the effect of external perturbations (Burdet et al., 2001; De Serres & Milner, 1991; Latash, 1992), stabilizing the limb during ball catching (Lacquaniti & Maioli, 1989) and improving accuracy of arm movment control (Gribble et al., 2003; Selen et al., 2006, 2009).

Despite the generally acknowledged role in impedance control and the recent interest in the mechanisms of recruitment of motor units in antagonist muscles during different tasks (Haynes & Kim, 2021), how the CNS regulates co-contraction is still unclear and two main hypotheses have been proposed. According to one hypothesis, during a co-contraction task (e.g., stiffening the arm by co-activation of antagonist muscles, Fig. 1A), the same synaptic inputs that drive motor neurons to generate endpoint force (Fig. 1B, *red* and *cyan* arrows) also modulate impedance (Berret & Jean, 2020; Forster et al., 2004; Hughes et al., 1995). Thus, the motor neuron pool of each co-active muscle receives a separate input both for stiffening (Fig. 1C) and force generation (Fig. 1D). In contrast, according to the second hypothesis, co-contraction is regulated through an additional separate synaptic input shared by the motor neuron pools of antagonist muscles (Fig. 1E-F) and therefore independent from the inputs underlying force generation (Borzelli et al., 2018; De Luca & Mambrito, 1987; Latash, 1992; Takagi et al., 2020).

**Figure 1:**
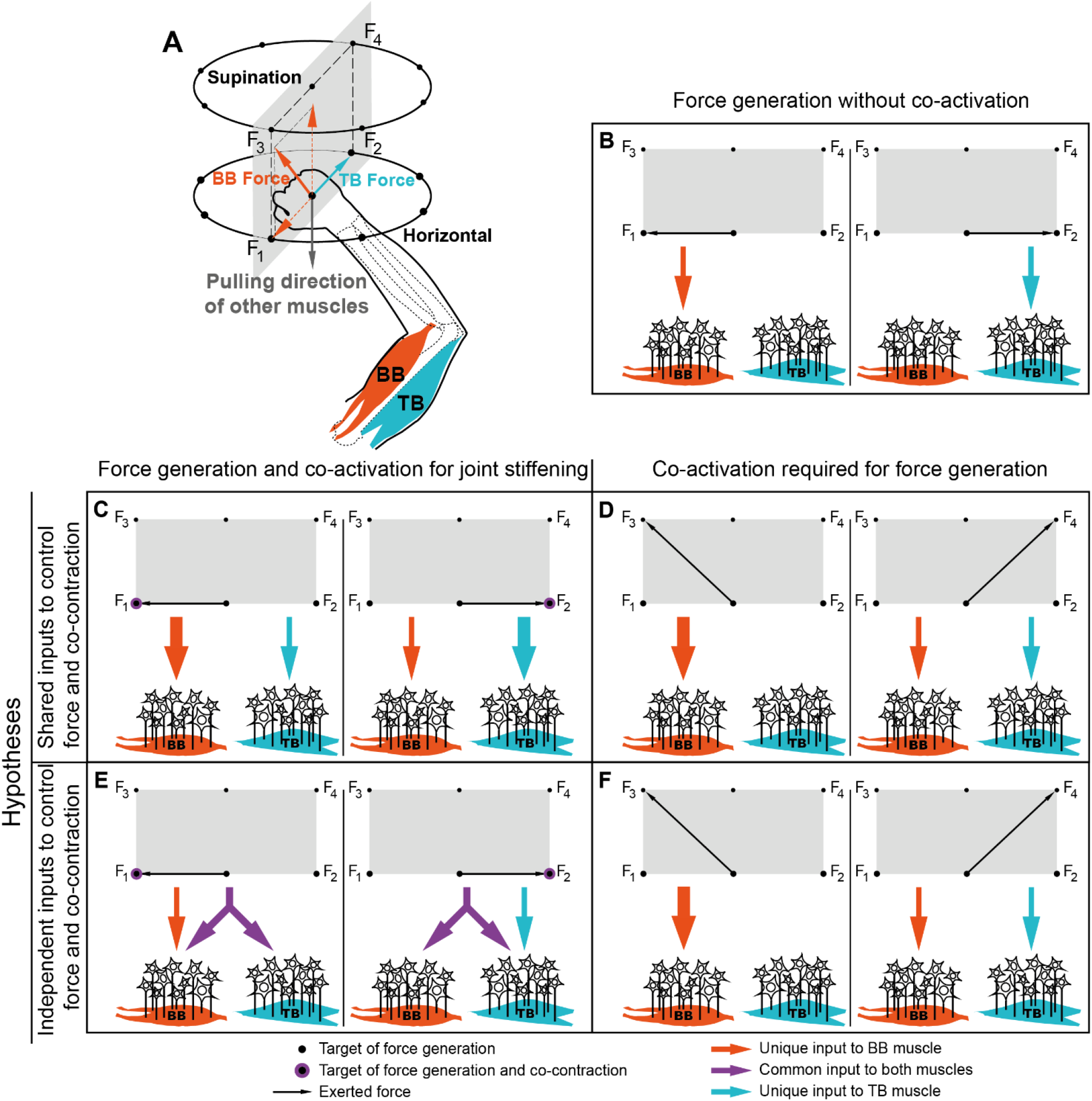
Conceptual models of synaptic inputs to motor neurons for the generation of force and the modulation of co-contraction. A. An upper-limb task designed to dissociate co-activation of antagonist muscles for impedance modulation and force generation. Force targets (black dots) arranged in two parallel planes can be reached by applying different combinations of torques at the upper-limb joints. Horizontal targets in the lower plane can be reached by different combinations of shoulder and elbow flexion or extension torques, producing horizontal forces at the hand. Supination targets in the higher plane requires the combination of shoulder and elbow torques (horizontal force) with wrist supination torque (producing a torque at the hand mapped as vertical displacement in this task). Two muscles are considered: Triceps Brachii (TB), an elbow extensor, i.e., with a pulling vector (cyan arrow) in the Horizontal plane (towards target F_2_), and Biceps Brachii (BB), whose action is both elbow flexion and forearm supination, i.e., with a pulling vector (red arrow) that has both a component (dashed arrows) in the Horizontal plane (towards target F_1_, opposite to TB) and a vertical component, required to reach the Supination plane. Thus, the co-activation of BB and TB may result in two actions, depending on the activation of other muscles (gray downward arrow in panel A): the increase of the elbow impedance through co-contraction with zero resultant force, or the simultaneous exertion of an endpoint force and supination torque. Four targets are highlighted: F_1_ and F_2_, lying on the Horizontal plane and requiring, respectively, an elbow flexion or an elbow extension torque to be reached, and F_3_ and F_4_, lying on the Supination plane and requiring wrist supination torque combined with, respectively, an elbow flexion or an elbow extension torque. B. Targets F_1_ or F_2_ requires separate synaptic inputs to MN pools of BB (red arrow) or TB (cyan arrow). C-D-E-F. Synaptic inputs to MN pools of BB and TB during different co-activation conditions according to the hypotheses of shared (panels C and D) or independent (panels E and F) inputs to control force and co-contraction. The hypothesis of shared control of force and co-contraction would predict the observation of independent synaptic input (red and cyan arrows) during co-activation both for force generation to reach Horizontal targets with co-contraction modulation (panel C) and for force generation to reach Supination targets without co-contraction modulation (panel D). On the contrary, the hypothesis of independent control of force and co-contraction would predict the observation of a shared common input (violet arrows) during Horizontal force generation with co-contraction modulation (panel E) but not during Supination force generation without co-contraction modulation (panel F).

It is well established that the common synaptic input within a given motoneuron pool, which determines the generated muscle force (Farina & Negro, 2015), can be identified using coherence analysis on the trains of action potentials of pairs of motoneurons (MNs) (Farmer et al., 1993). Recently, coherence analysis has also been used to assess the synchronous modulation of the neural drive to MNs across muscles, identified from High-Density EMGs (Holobar & Zazula, 2007), and to thus unveil synaptic inputs shared across muscles (Del Vecchio et al., 2019, 2022; Laine et al., 2015; Martinez-Valdes et al., 2018).

In this study, we first identified the spike trains of MNs by decomposing High-Density surface electromyograms (HD EMGs) recorded from Biceps Brachii (BB) and Triceps Brachii (TB) muscles during the performance of a task requiring to concurrently generate multi-directional isometric forces and modulate co-contraction. We then performed an analysis of coherence of the spike trains to assess the existence of a shared input to the MNs of antagonist muscles. The presence of a significant peak in the coherence between the discharge trains of the pools of MNs of antagonist muscles during a co-contraction task, together with the absence of the peak during force generation without co-contraction (see Fig. 1B and F), revealed a shared inputs to these muscles (Fig. 1E) and supported the existence of a specific input controlling the modulation of co-contraction independently from the generation of force. The occurrence of a peak in the beta band, identified across the generation of forces along different directions, suggested a cortical orgin of such shared input (Farina & Negro, 2015). We also assessed the coherence of the spike trains of MNs of each muscle after removing the common input to the antagonist muscle or the exerted force. Such residual coherence (Laine et al., 2015) allows to distinguish the MNs recruited to modulate co-contraction from those involved in force generation. The identification of MNs selectively driven to control force or co-contraction suggests the existence of specific neural patways. To the best of our knowledge, our results provide, for the first time, evidence for the existence of a synaptic input to motoneurons that specifically controls co-contraction, and that is independent from the input driving the generation of force.

## Methods

Eight healthy male adults (age range: 22 - 35 years) participated in the study after giving written consent.

All participants had normal or corrected to normal vision and did not report having had any neurological disorder or upper right limb injury. All procedures were conducted in accordance with the Declaration of Helsinki and were approved by the ethics committee IRCCS Sicilia-Sezione Neurolesi “Bonino-Pulejo” (Prot. n. 02/18).

### Setup

Participants sat on a gaming chair in front of a desktop with car safety belts immobilizing their torso and shoulders (see Figure 2A). The right hand was inserted in an orthosis, rigidly connected to a cylindrical handle that participants could grasp to easily generate supination torques, and to a 6-axis force transducer (Delta F/T Sensor, ATI Industrial Automation, Apex, NC, USA). The height of the chair and its distance from the table were set to ensure the elbow angle was 90° (0° full extension) and the hand was at the height of the solar plexus. A horizontal mirror, preventing the view of the participant’s hand, reflected the image displayed by a monitor rendered at 120 Hz (60 Hz per eye). Participants wore shutter glasses (GeForce 3D Vision 2, NVIDIA Corporation, Santa Clara, CA, USA) that allowed the stereoscopic vision of a three-dimensional scene. The scene comprised a desktop and a cursor whose displacement was computed in real time from the measured isometric force and torque and from surface EMGs collected with bipolar electrodes on multiple muscles acting on the elbow and shoulder joints (see Paragraph ‘Bipolar surface electromyography ‘, below). At rest, and without perturbing forces, the cursor was displayed at a position corresponding to the center of the palm (rest position). The displacement in the horizontal plane was proportional to the force generated in the horizontal plane, while the vertical displacement was proportional to the supination torque.

**Figure 2:**
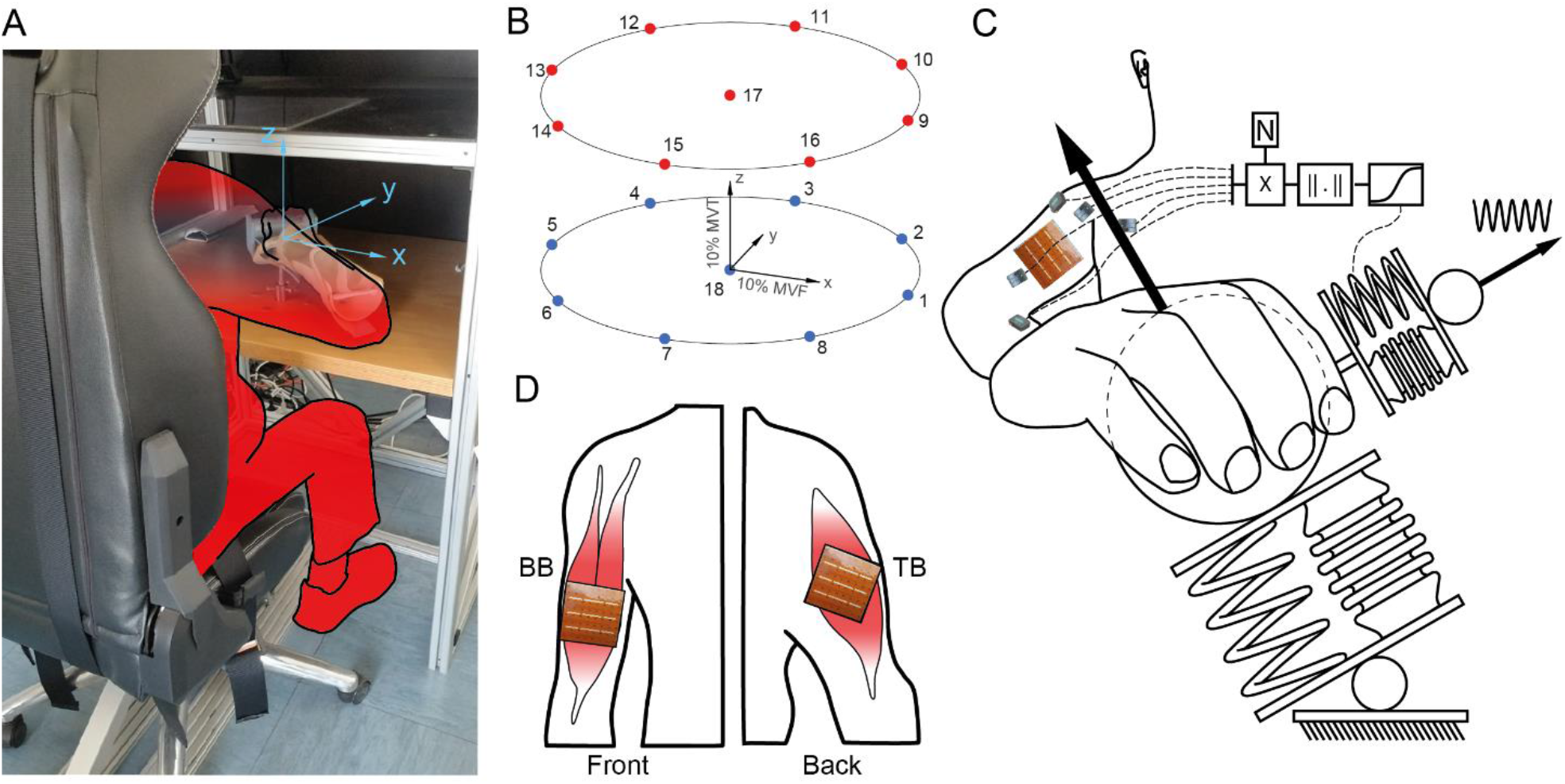
Experimental setup. A. Participants inserted their right forearm into an orthosis rigidly connected to a 6-axis force transducer, fixed below a desktop. Participant viewed a virtual scene that consisted of a spherical cursor whose displacement from the rest position (i.e. the position that the cursor assumed when the muscles were relaxed) was proportional to the exerted force and torque. The origin of the coordinate system used to displace the cursor in the virtual scene was placed at the center of the participant’s palm. Participants could move the cursor in the horizontal plane by exerting horizontal forces (x-axis: medio-lateral direction; y-axis: dorso-ventral direction), and along the vertical direction (z-axis: upward vertical direction) exerting a forearm supination torque. B. Participants were instructed to displace the cursor to reach one of 18 spherical targets arranged in two parallel horizontal (x-y) planes. Horizontal targets (blue) lay in the lower horizontal plane and Supination targets (red) lay in the upper horizontal plane and required a supination torque to be reached. C. During Perturbation blocks, the motion of the cursor was simulated as two mass-spring-damper systems (MSDs) in series. The first MSD was displaced by the force exerted by the participant, recorded by the force transducer, and determined the mean position of the cursor. The second MSD was perturbed by a simulated external unpredictable force and determined the oscillations around the mean position. Participant could reduce the magnitude of the oscillations of the second MSD system in real-time by voluntarily co-contracting their muscles. D. High-density EMG grids covered the BB (left) and TB lateral head (right) muscles.

### Experimental design

After an initial familiarization phase, participants performed 3 blocks of trials. In the first block, denoted Maximal Voluntary Force and Torque (MVFT) block, they were asked to exert their Maximum Voluntary Force (MVF) and Maximum Voluntary Torque (MVT). The MVF was the maximum force that participants could generate towards their chest along the horizontal plane (-y direction; Figure 2A) and the MVT was the maximum torque that participants could generate supinating their wrist (+z direction). Participants alternated two repetitions of MVF with two repetitions of MVT and the peak values were used to normalize the target positions in the following blocks (MVF normalized the position along the horizontal x-y plane, MVT normalized the target position along the vertical z axis). The second block (Baseline) was composed of 17 trials during which participants were asked to move a spherical cursor from the rest position to a target position, located in one of 17 spatial positions (see Figure 2B). Similarly, the third block (Perturbation) was composed of 18 trials during which participants were asked to stabilize the cursor in the rest position by increasing muscular co-contraction, or to move it from the rest position to a target position, located in one of the 17 spatial positions also used in the Baseline block. At the beginning of each trial (rest phase), participants were asked to relax their right arm muscles to maintain the cursor inside a semi-transparent sphere in the rest position. After doing so for 1 s, the sphere disappeared and reappeared in one of the target positions (target go event). One subset of targets included eight targets equally spaced along a circumference in the horizontal plane with 20% MVF radius (targets 1 to 8 in Figure 2B), together with the target in the rest position (target 18, only in Perturbation block). No supination torque was required to reach this set of *horizontal targets.* The other subset included 9 targets at the same horizontal positions but with an offset of 20% MVT in the +z direction (targets 9-17 in Figure 2B). These *supination targets* could be reached only by eliciting a 20% MVT supination torque.

Participants were asked to displace the cursor to reach the target and to maintain the cursor within the target sphere, whose radius exceeded that of the cursor by 4% MVFT, for 20 s (holding phase). Targets were displayed in random order and breaks of 40 s were inserted between trials to avoid fatigue. Bipolar EMGs and force data collected during the holding phase of the Baseline block were used to estimate a subject-specific matrix that approximates the mapping of EMG amplitude onto isometric force (EMG-to-force matrix, see below) and its null space. The maximum amplitude of each rectified and low-pass filtered (second-order Butterworth; 1 Hz cutoff) EMG collected during the same phase was used to normalize EMGs during the Perturbation block. High-density EMGs obtained during the hold phase of all trials of both Baseline and Perturbation blocks were processed for the identification of motor units (MUs). While during the Baseline block the motion of the cursor was proportional to the exerted force and torque and it was simulated as a single adaptive mass-spring-damper system (MSD), during the Perturbation block it was simulated as two MSDs in series (see Figure 2C), as described in detail in (Borzelli et al., 2018). Briefly, the force generated by the subject was applied on the first MSD while a simulated sinusoidal force, applied to the second MSD, perturbed the motion of the cursor. The stiffness of the second MSD was modulated in real-time according to the norm of the projection of the muscle activation vector component along the null space of the EMG-to-force matrix, through a logistic function. Therefore, during the Perturbation block, participants could reduce the oscillation of the cursor by co-contracting shoulder and elbow muscles, without affecting the generated endpoint force required to reach the target.

Experiment control, data acquisition (force, torque, and bipolar EMG), and data analysis were performed with custom-written software in MATLAB^®^ (MathWorks Inc., Natick, MA) and Java^®^.

### High-density surface electromyography

Surface EMGs of both heads of Biceps Brachii (BB) and the lateral head of Triceps Brachii (TB, see Figure 2D) of each participant were recorded with two arrays, each with 8 x 8 electrodes (3 mm diameter, 10 mm interelectrode distance; HD10MM0808; OT Bioelettronica, Turin, Italy). Participants had their skin cleansed with abrasive paste and the arrays were centered between the proximal and distal tendons of the muscles, with rows visually aligned parallel to the muscle fibers. The array on the BB was placed such that the junction between the long and short heads, identified by palpation, was located between columns 4 and 5. The TB array covered the muscle belly, identified by palpation, and the columns were parallel to the muscle fibers, as indicated by recommendations from SENIAM (Hermens et al., 1999) and Kendall (Kendall et al., 1994). Monopolar EMGs, referenced to the wrist, were amplified and recorded (2048 Hz sampling rate) using a 16-bit A/D converter (Quattrocento multi-channel amplifier; OT Bioelettronica, Turin, Italy).

### Bipolar surface electromyography

Bipolar, surface EMGs were recorded from eleven muscles crossing the shoulder and elbow joints: brachioradialis, biceps brachii, pectoralis major, anterior deltoid, middle deltoid, posterior deltoid, triceps brachii long head, infraspinatus, teres major, latissimus dorsi, and middle trapezius. Participants’ skin was cleaned with alcohol and electrodes were placed based on recommendations from SENIAM (Hermens et al., 1999) and by palpation to locate the muscle belly and visually orienting the pair of electrodes along the expected direction of fibers. The single bipolar electrode on BB, which was required for the real-time estimation of the null space component of the muscle activation (see Paragraph ‘Experimental design’, above), was placed just distally to the high-density grid and aligned parallel to the BB long head longitudinal axis. Owing to the skin-parallel fibered architecture of BB, bipolar electrodes placed just distally to the grid would be expected to provide EMGs as representative as those detected more proximally and not in proximity to the muscle innervation zone (Vieira & Botter, 2021). Bipolar signals were acquired at 1000 Hz with active wireless bipolar surface electrodes (Trigno System, Delsys Inc., Natick, MA, USA), bandpass filtered (20 – 450 Hz), and amplified with a 1000 gain. High-density and bipolar EMGs and force data were recorded by two different data acquisition systems and were synchronized offline using a trigger signal delivered at the beginning of each acquisition by one system and recorded by the second system.

### EMG-to-force mapping

Isometric generation of submaximal force and torque allowed to use a linear approximation of the relation between EMG amplitude and force and torque:

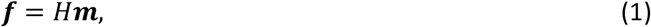

where ***f*** is a tridimensional vector composed by two components of force exerted along the horizontal plane and one component of supination torque, ***m*** is a 11-dimesional muscle activation vector, and *H* is an EMG-to-force matrix that maps muscles activation onto force. The matrix *H* was estimated using multiple linear regressions of each low-pass filtered (second-order Butterworth; 1 Hz cutoff) force component, on the rectified, low-pass filtered (second-order Butterworth; 1 Hz cutoff) and normalized to the MVFT bipolar EMGs recorded during the holding phase of the Baseline block. The matrix *H* was also used to compute a null space matrix *N*, i.e. an orthonormal basis spanning the subspace of vectors *n* that are mapped by the H matrix onto the null force vector:

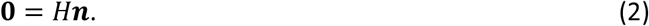

### Motor unit decomposition

Monopolar, high-density EMGs were filtered (2nd order Butterworth, 15-350 Hz) and decomposed through an automatic and validated algorithm (Holobar & Zazula, 2007), separately for each 20-s holding phase. After decomposition, the firing instants of identified MUs were used to trigger and average single differential EMGs over 30 ms epochs and inspected for spurious units (Del Vecchio et al., 2020; Power et al., 2022). Only units whose firing rates were stable over the entire contraction (pauses shorter than 500 ms) and whose action potential propagation (i.e. the shifting in time of the action potential peak, identified on channels placed along the fiber direction, proportionally to its spatial distance from the innervation zone) could be appreciated, were used in analysis. Moreover, mean discharge rates of MUs identified on the BB were required to fall between 7 and 51 Hz and mean discharge rates of MUs identified on the TB were required to fall between 15 and 37 Hz (Enoka & Fuglevand, 2001). The activity of each MU was expressed as a binary spike train in which each time sample (2048 Hz sampling frequency) was assigned either a value of 1 if it marked the beginning of an action potential (identified by the algorithm for MU decomposition) or a value of 0 otherwise. Trials in which fewer than three motor units were decomposed were excluded from the following analyses. In the following analysis, we replicated, on the BB and the TB muscles, the approach that Laine and collaborators (2015) proposed to determine the cross-muscle coherence, the total within muscle coherence, and the residual within muscle coherence on the Vastus Medialis and Vastus Lateralis muscles.

### Cross-muscle analysis

We tested the synchronous modulation of the firing rates of motor unit pairs using coherence analysis. Coherence (Rosenberg et al., 1989) is a frequency domain extension of Pearson’s correlation, estimating the linear correlation between the two signals at any given frequency.

In the cross-muscle analysis (see Figure 3), we calculated the coherence between the spike trains of all pairs of MUs identified in the two antagonist muscles. As proposed by Laine and collaborators (2015), all unique pairs of spike trains, identified on the BB and on the TB muscles during a trial, were concatenated into two long trains, which were then subjected to coherence analysis. Each MU train of spikes (here called *i* and *j*) was divided into segments of 3 s length, and the FFT of each segment *u* (*I_u_* and *J_u_* respectively) was calculated considering a 0% overlapping and weighted by a rectangular window function. The complex values obtained across the segments at each frequency were used to derive the auto-spectra (*ii*, and *jj*) and cross-spectra (*ij*) of *i* and *j* spike trains:

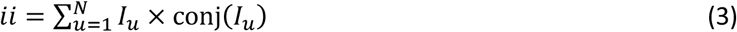

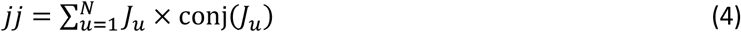

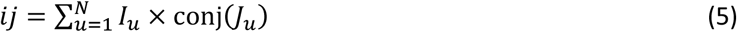

Where conj() refers to the complex conjugate of *I_u_* and *J_u_* and *N* = 7 is the number of segments of 2.8 s length.

**Figure 3:**
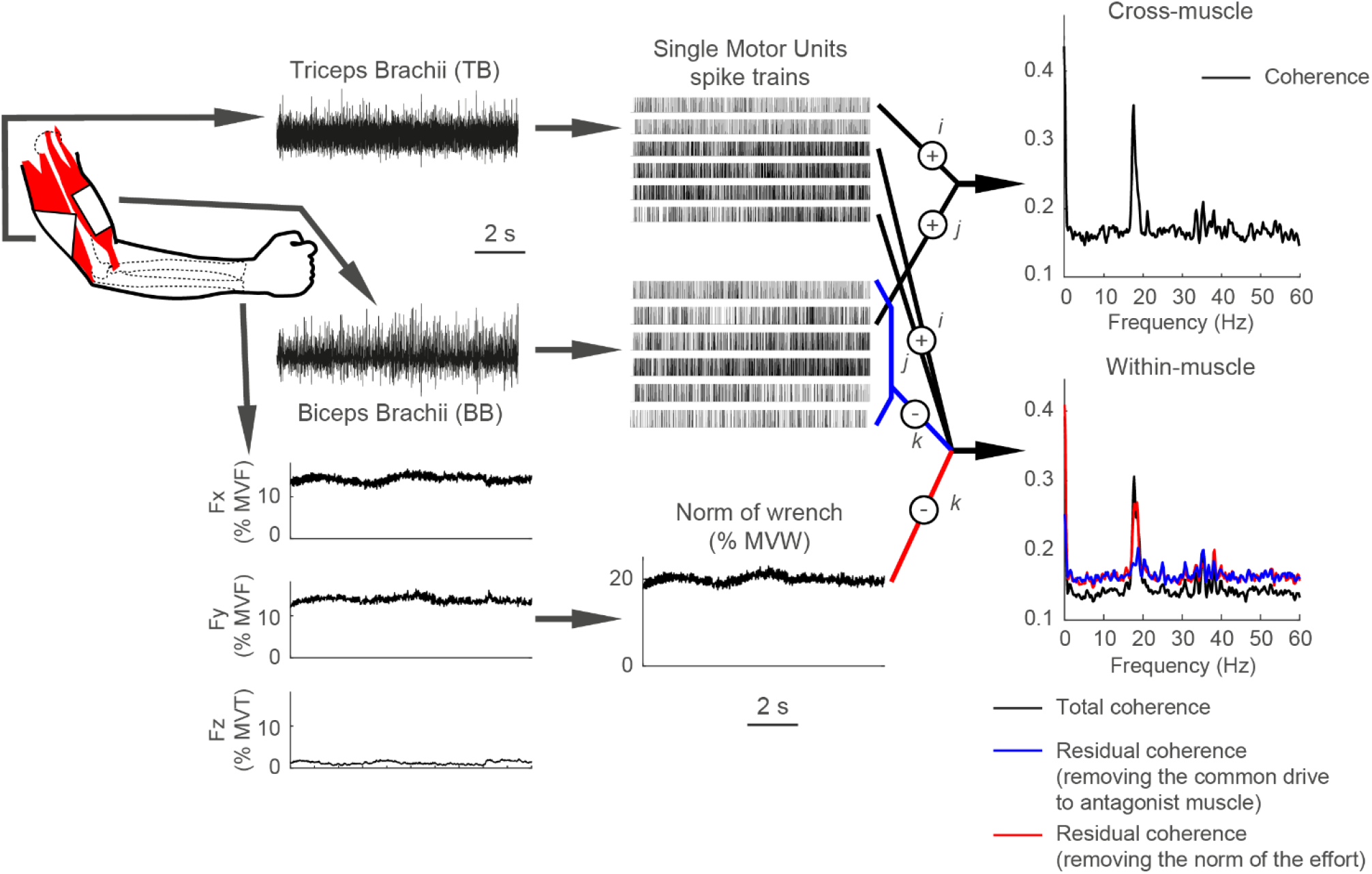
Recordings of high-density EMG signal and data analysis. High-density EMG signals (64 channels per muscle) were recorded from TB and BB during submaximal, isometric contractions. These signals were decomposed to reveal the firing pattern of single MUs. Two approaches were tested. In a cross-muscle analysis (top right panel), we calculated the coherence for paired BB-TB MUs (i and j signals in Equations 3–6) identified during a single trial (the example shows data collected during the generation of force to reach target 2 of the Perturbation block, see Figure 2B). In a within-muscle analysis (bottom right panel), we calculated the total coherence between all pairs of MUs from the same TB muscle (i and j in Equations 3–6; black trace). We further computed the residual coherence, which represents the within-muscle MU coherence after removing the common neural drive to the antagonist muscle (blue trace) or the effect of force/torque modulation (red trace) contribution (k in Equations 7–14).

The magnitude squared coherence (typically referred to as “coherence”) for each frequency was then calculated as:

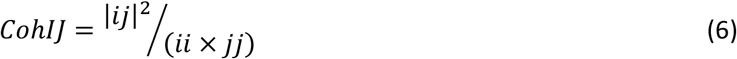

Each coherence profile was then smoothed in the frequency domain using a three-point running median window.

Co-activation of BB and TB is required both to modulate the elbow impedance (co-contraction, i.e. without the generation of end-point force and torque) or to move the cursor toward a target (e.g. target 2 in Figure 2B), which required the activation of the TB to move the cursor along the horizontal plane, and the activation of BB to generate the supination torque required to move the cursor along the vertical direction. To distinguish these two conditions, the cross-muscle analysis was performed separately on two subsets of target: horizontal targets during the Perturbation block, in which the co-activation was due to the increase in elbow impedance, and supination targets during the Baseline block, in which no impedance modulation was required, and therefore co-activation was required to reach the target.

### Within-muscle analysis

In the within-muscle analysis, we isolated the neural drive to each muscle from the contribution of any common drive shared between muscles and responsible for the modulation of force and torque. More specifically, as proposed by Laine and collaborators (2015), we used Equations (3–6) to compute the total coherence, with *I_u_* and *J_u_* corresponding to the FFT of all concatenated unique pairs of spike trains identified on the same muscle. The residual coherence is an estimation of the synchronous modulation of the firing rate of two MUs (*i* and *j)* in the same muscle, after removing the component that is synchronous to a third reference signal (*k*). In this analysis *k* represented the common neural activity of all units of the antagonist muscle, calculated as the summation of all individual MU spike trains recorded from the antagonist muscle (Farina, Negro, et al., 2014; Laine et al., 2015), or as the norm of the force and torque, normalized to the MVFT (Figure 3).

We first derived the auto-spectra *kk* for the reference signal *k* as described for *i* and *j* above (3-4). Then, the cross-spectra of signal *k* with signals *i* and *j* were calculated as below:

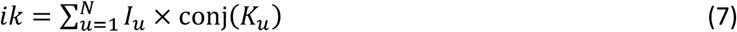

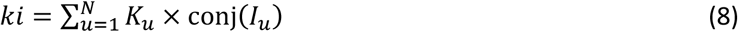

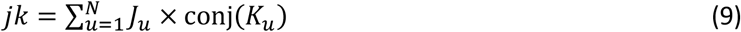

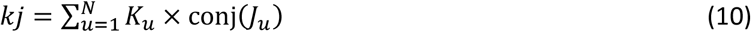

And the cross-spectra and auto-spectra between *i* and *j* not due to *k* signal were calculated as follows:

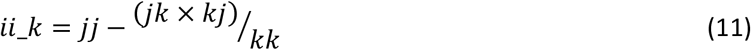

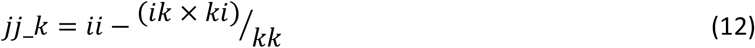

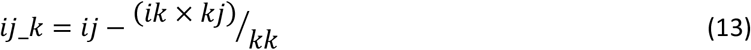

Therefore, the residual coherence between *i* and *j* after removing the component synchronized with *k* was calculated as follows:

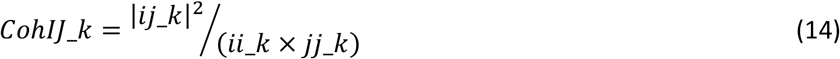

Residual coherence was computed and then averaged for all pairs of units (*i* and *j)* in each of the two muscles, one at a time, with the sum of the spike trains of the other muscle or the norm of the force and torque (normalized to the MVFT), providing the reference signal (*k*). Total coherence, residual coherence after removing the component synchronous to the common drive to the antagonist muscle, and residual coherence after removing the component synchronous to the norm of the force and torque were computed separately for each participant and trial of the Perturbation block.

### Task-based separation of motor units

A significant total coherence and a significant residual coherence in the same frequency band in a MU pair, identified during the same trial on the same muscle, indicates that the two units synchronously modulate their firing pattern at that frequency without sharing a common input with the reference signal. On the contrary, a significant total coherence without a significant residual coherence indicates that the two units share the same input with the reference signal. Therefore, a pair of MUs was considered to be driven by a common synaptic input specific for force generation if they showed a significant total coherence and a significant residual coherence when excluding the component synchronized with the common drive to the antagonist muscle, but did not show a significant residual coherence when excluding the component synchronized with the force and torque in the [12 21] Hz band (see Results for the definition of this band). Similarly, a pair of MUs was considered to be driven by a common synaptic input specific for co-contraction modulation if they showed a significant total coherence and a significant residual coherence when excluding the component synchronized with force and torque but did not show a significant residual coherence when excluding the component synchronized with the common drive to the antagonist muscle in the [13 21] Hz band. Therefore, we defined two subsets of MUs, one recruited only to generate force and the other recruited only to modulate co-contraction, as those MUs that satisfied, in all pairings with other units extracted from the same muscle during the same trial, respectively only the criteria for force generation or for co-contraction modulation. The approach proposed for the assignation of a MU to a subset is strict, because the violation of the criterion for that subset, even at a single frequency bin (in the [13 21] Hz band) during a single pairing would exclude that MU from the selected subset. For this analysis each MU train of spikes was divided into segments of 1 s length.

Finally, we calculated the total coherence among all pairs of MUs, identified on the same trial, within these subsets of MUs to determine the frequency band of their common neural input.

### Statistics

Based on the null hypothesis that no coherence exists at a given frequency, we defined a 95% confidence level (Laine et al., 2015). For each coherence profile, the 95% confidence level (*CL_TOT_*) was derived (Carter, 1987; Rosenberg et al., 1998) as:

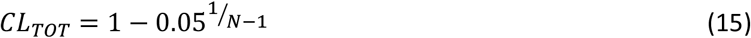

Where *N* is the number of data segments used to calculate the coherence profile.

For each residual coherence profile, the 95% confidence level (*CL_RES_*) was derived (Carter, 1987; Rosenberg et al., 1998) as:

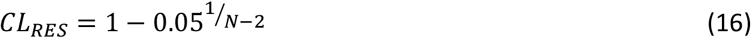

Then, a binomial test was used to determine whether the proportion of pairs of units identified for all trials and participants, showing significant coherence, exceeded the rate expected by chance. The test provided a conservative evaluation of the relevant frequency content of neural drive within and between muscles.

## Results

### Motor unit decomposition

After exclusion of MUs with a non-physiological action potential or firing rate and trials in which no MUs were identified on the BB or on the TB (Figure 4), a total of 896 MUs [median (interquartile interval) across subjects 126 (89.5)], identified during the Baseline block [N = 265 units, 34 (51)] and the Perturbation block [N = 631 units; 78 (71)], were retained for analysis. Of these, the number of units identified for BB during the Baseline block [N = 175 units; 20.5 (35)] was about half of the number obtained during the Perturbation block [N = 309 units; 38.5 (44)], while the number of TB MUs identified during Baseline [N = 90 units; 7.5 (17)] was less than one third of the number obtained during the Perturbation block [N = 322 units; 33.5 (31)]. The MU discharge rates during the baseline block were 25.2 ± 9.6 (mean ± std) for the BB and 21.6 ± 5.3 for the TB, and during the perturbed block were 20.4 ± 8.7 for the BB and 19.8 ± 4.3 for the TB, in line with literature (Freund, 1983). In Figure 4, examples of data collected from BB and TB during the exertion of two force target of the baseline block, and the related MUs extracted from data, are shown.

**Figure 4:**
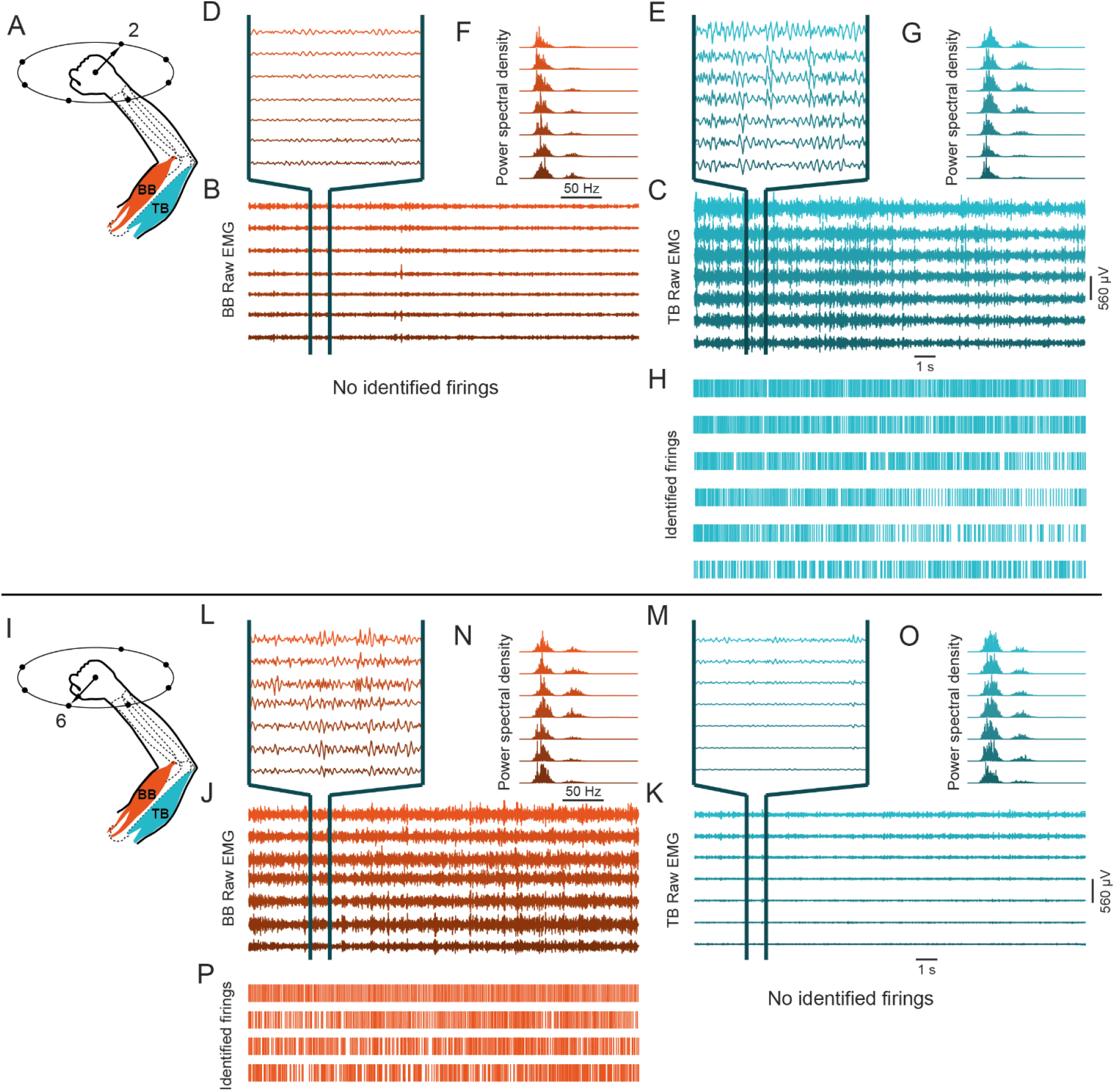
Example of the EMG signal acquired by a column of channels collecting the activity of BB and another collecting the activity of TB and the identified firings. The data were collected during the generation of force along two opposite directions on the horizontal plane: one, along target 2 (see Figure 1B), required a torque of elbow extension (A) while the other, along target 6, required a torque of elbow flexion (I). The raw (unprocessed) EMG activity collected from the BB (panels B and J, with a portion of the signals expanded respectively in panels D and L) and from the TB (panels C and K, expanded in panels E and M) shows a typical power spectral density (panels F, g, N, and O). The firings of the identified MUs on the TB during the generation of a force along target 2 are reported in panel H, while the firings of the identified MUs on the BB during the generation of a force along target 6 are reported in panel P. Consistently with the observation that BB was inactive during the generation of a force along target 2 and TB was inactive during the generation of a force along target 6, no MUs were identified.

After exclusion of any trials in which fewer than three motor units were decomposed, a total dataset of 35 trials of the baseline block, lying on the supination plane, and 38 trials of the perturbed block, lying on the horizontal plane, from across eight subjects was available for cross-muscle coherence analysis. A total dataset of 73 and 71 trials of perturbed block was available for within-muscle coherence analysis on TB and BB respectively.

### Cross-muscle analysis

Figure 5 depicts the percentage of trials MUs that showed significant BB-TB coherence (95% confidence level) coherence identified during the generation of force to reach the horizontal targets of the Perturbation block (Figure 5A) or the horizontal targets of the supination targets of the Baseline block (Figure 5B). Fractions greater than the marked confidence level (dashed line) indicated that significant coherence occurred in a substantially higher number of trials than expected by chance.

**Figure 5:**
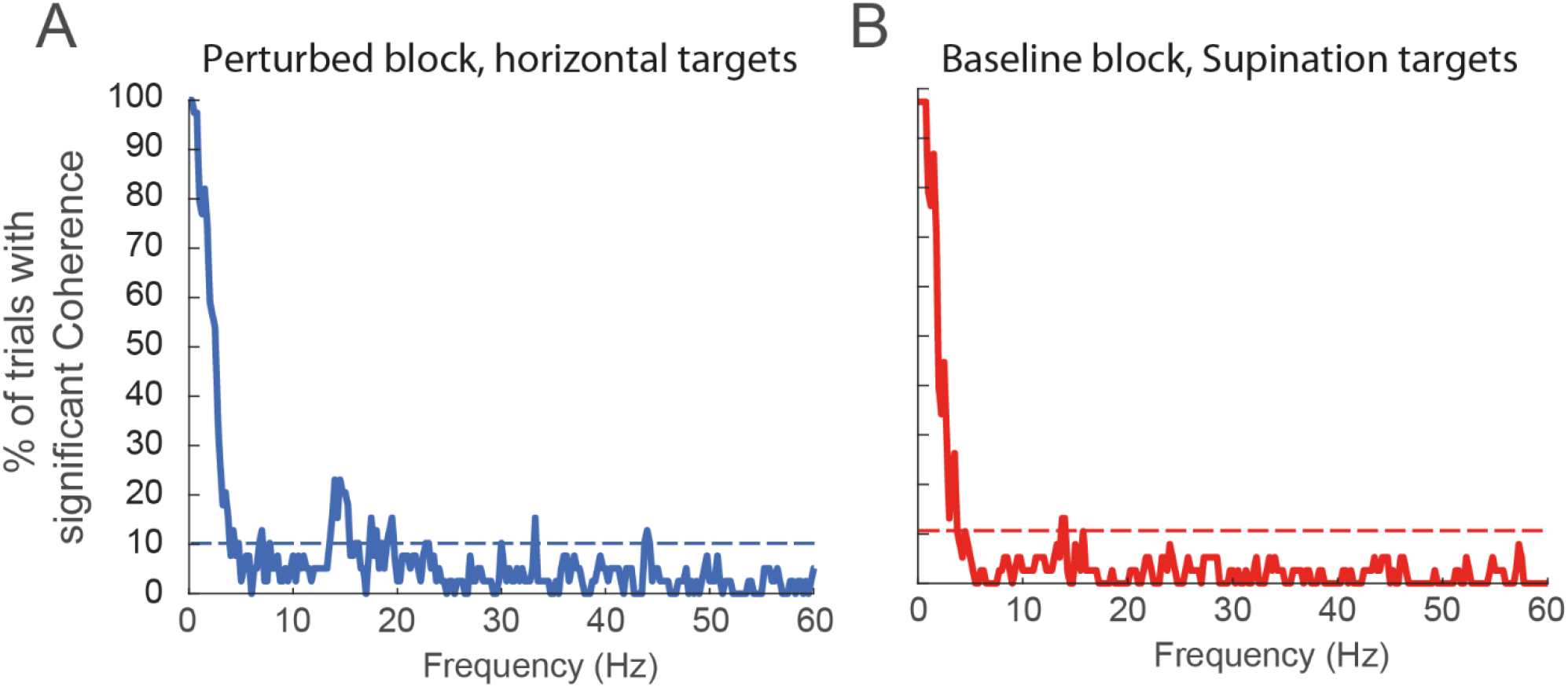
Cross-muscle analysis: percentage of the of the pairs of MUs, identified on antagonist muscles, showing a significant coherence. The coherence analysis was performed on all pairs of MUs, identified on the BB and the TB muscles during the horizontal targets of the Perturbation block (A) and the supination targets of the Baseline block (B). The dashed horizontal line indicates the highest proportion that could have been observed simply by chance (95% confidence level). The threshold indicating the higher percentage of trials that may show significant coherence by chance (95% confidence level) is indicated by the horizontal dashed lines.

The cross-muscle coherence analysis performed on both the target sets, identified a [0 5] Hz component of the synaptic input shared between the muscles and another component at around 15 Hz, that was more marked when muscles were recruited to generate force along the horizontal plane during the perturbed block. Remarkably, another significant peak was identified at around 20 Hz only when participants were required to exert a force and to increase BB-TB co-contraction (Figure 5A). This narrow peaks at around 20 Hz and the wider peak at 15 Hz in the coherence between the discharge trains of the pools of MNs located on antagonist muscles, identified during tasks requiring co-contraction for impedance modulation (horizional targe targets in the pertuebed block) revealed the existence of shared inputs to these muscles and supported the hypothesis of a specific input for the modulation of co-contraction that is independent from the input for the generation of force.

### Task-based separation of motor units

While the cross-muscle analysis investigated the coherence between pairs of units identified on antagonist muscles, the within-muscle analysis investigated the coherence between units identified on the same muscle, considering all frequency components (total coherence) or excluding the components synchronized with the common drive to the antagonist muscle or with the exerted force and torque (residual coherence). Figure 6A and B depicts the percentage of MU pairs, identified on all trials of the Perturbation block, that showed a significant total within-muscle coherence, for TB (Figure 6A) and BB (Figure 6B). Besides the <5Hz significant peak, TB showed frequency bins with a significant percentage (i.e. above the dashed horizontal line) of pairs of MUs with total coherence in the 12 to 21 Hz band and some peaks at around 30Hz. Differently from the TB, the total coherence for the BB muscle showed, besides the <5Hz peak, only some significant coherence in the 10 to 15 Hz band.

**Figure 6:**
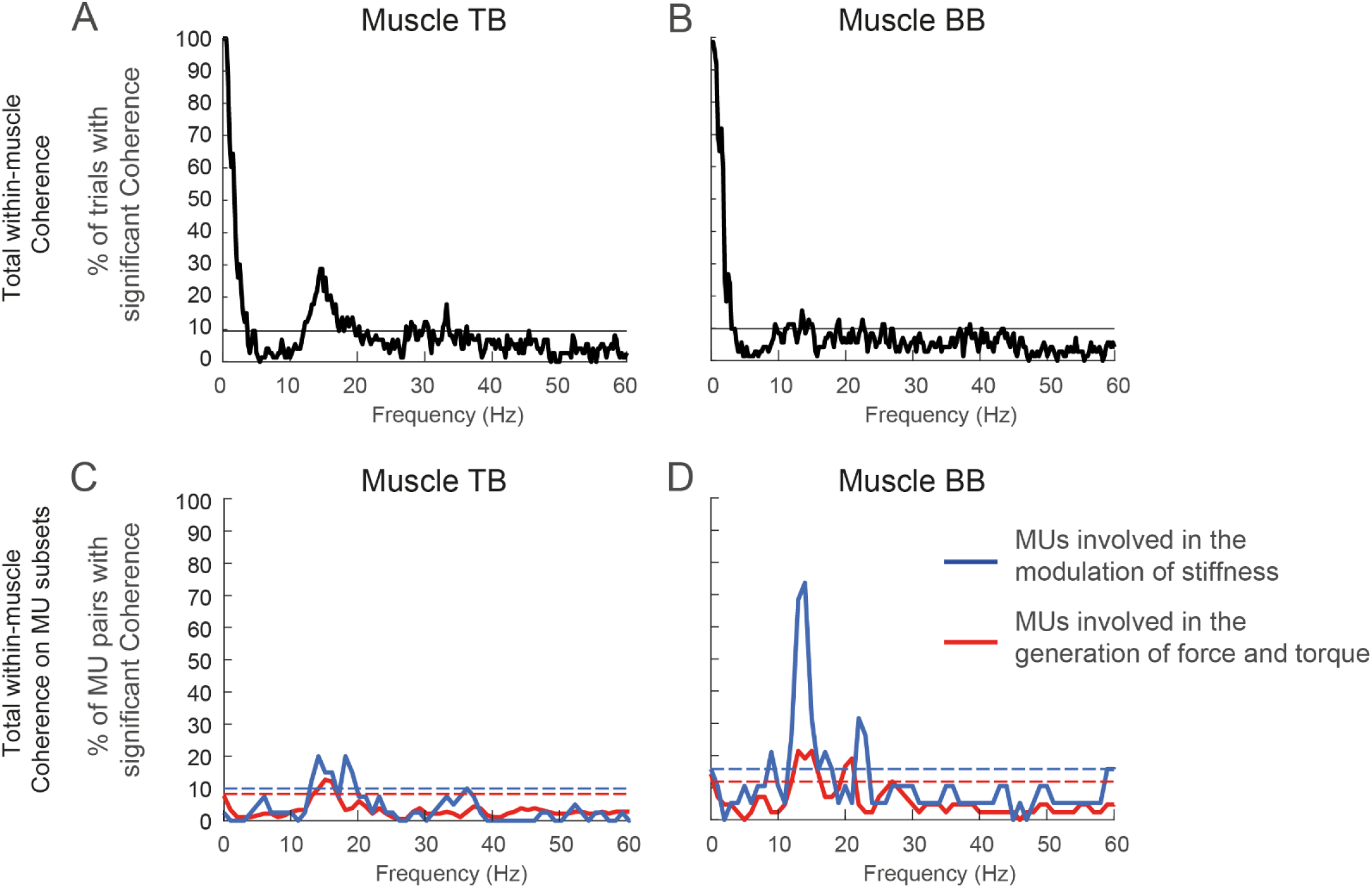
Total within-muscle MU coherence and common input to MUs driving the generation of force and torque or the modulation of co-contraction. Top panels represent the percentage of the pair of MUs, identified on the TB (A) or on the BB (B), that showed a significant total within-muscle coherence. Bottom panels represent the total within-muscle coherence separately calculated among the subset of MUs selectively recruited to modulate co-contraction or generate force and torque in TB (C) and BB (D). The fraction of pairs that showed a significant coherence is displayed in the figure (blue: MUs selectively recruited to modulate co-contraction; red MUs selectively recruited to generate force and torque). In all panels, the horizontal dashed lines represent the threshold for significance, i.e. the higher value of the 95% confidence interval of the percentage of MU pairs that may show significant coherence by chance. In panels C and D, the difference between the confidence levels is due to the different numbers of pairs of MUs involved in the modulation of co-contraction or in the generation of force and torque.

We then used the total and residual within-muscle coherence of to identify two subsets of MUs in each muscle. A MU, recorded on the TB or on the BB, was identified as recruited selectively to perform a task, according to its total and residual coherence in the 12 to 21 Hz band, wherein the total within-muscle coherence exceeded the chance threshold. During the Perturbation block, 9% of TB MUs and 8% of BB MUs were selectively recruited to modulate the co-contraction while 22% of TB MUs and 14% of BB MUs were selectively recruited to generate force, therefore providing 31% of TB units and 22% of BB units for analysis. The total coherence among these subsets of units revealed peaks at different frequency bands. While both MUs of the TB (see Figure 6C) contributing to force generation and to co-contraction showed a significant coherence peak at 15 Hz, only MUs contributing to co-contraction showed a second significant coherence peak at 20 Hz. On the contrary, both MUs of the BB (see Figure 6D) contributing to force generation and to co-contraction showed a significant coherence peaks at 15 Hz and another peak at higher frequencies, but, this second peak in the coherence within the MU pool contributing to co-contraction, occurred at higher frequencies (22 Hz) with respect to the coherence peak within the MU pool contributing to force (20 Hz).

## Discussion

We investigated the control of antagonist muscles at the motor neuron level and the existence of a neural pathway responsible for modulating limb impedance that is separate from the pathways underlying the generation of force. To disentangle the activation of a pair of muscles required for force generation and for impedance modulation we employed a recently developed approach, ‘virtual stiffness’ (Borzelli et al., 2018), to instruct participants to modulate co-contraction in several upper limb muscles, using real-time feedback of co-contraction level as stiffness of a virtual end-effector. The firings of MUs were identified by decomposing the high density EMG signal through an established algorithm (Holobar & Zazula, 2007) and the common drive to motor units of two antagonist muscles was identified through coherence analysis (De Luca & Erim, 1994). A cross-muscle analysis detected a significant coherence peak in the alpha band (close to 15 Hz) between the discharge trains of motor neurons pools of BB and TB that occurred in a larger fraction of trials requiring co-activation to modulate co-contraction. A second wide significant band (between 15 and 20 Hz) and other peaks (at 33 and 43 Hz) appeared at higher frequencies only when BB and TB were co-activated to modulate co-contraction, thus suggesting the existence of an independent synaptic input for modulating co-contraction. A within-muscle analysis identified subsets of MUs that were selectively recruited either to generate force or to modulate co-contraction. While a peak at 15 Hz was identified in BB and in TB both among MUs recruited to generate force or to modulate co-contraction, in TB a second higher peak appeared only within the set of MUs recruited to modulate co-contraction and in BB this second peak occurred within the set of MUs recruited to modulate co-contraction at higher frequencies (22 Hz) with respect to the set of MUs recruited to generate force (20 Hz).

The control strategies exploited by CNS to regulate limb impedance are still debated. While some studies have suggested an independent recruitment of individual muscles to optimally control both force and stiffness (Berret & Jean, 2020; Forster et al., 2004; Hughes et al., 1995) (shared inputs in Figure 1), other studies have proposed a separation of the control of force and stiffness, through shared common input to antagonist muscles (Borzelli et al., 2018; De Luca & Mambrito, 1987; Latash, 1992; Takagi et al., 2020) (independent inputs in Figure 1). To address this question, we took advantage of the “virtual stiffness” approach, which allowed the discrimination of the neural drives responsible for the co-activation for impedance modulation from the co-activation responsible for the generation of force in a direction that requires their summed contribution. Therefore, this is the first time the modulation of co-contraction for impedance generation is assessed at the MN levels. Recently the coherence between antagonist muscles during a co-contraction task was investigated with EMG (Nandi et al., 2019; Ohtsuka et al., 2022), which can be affected by cross-talk and decorrelation effects and whose interpretation is complicated by the fact that the EMG signal contains both information about the neural drive to the muscles and the electrical properties of the muscle fibers (Dideriksen et al., 2018; Farina, Merletti, et al., 2014).

Results presented in this study provide new insights on the physiological mechanism underlying the modulation of impedance at the MN level. Despite the synchronization between MUs of different muscles has long been established (Datta & Stephens, 1990; De Luca & Erim, 1994), its anatomical origin and physiological role is unknown. Significant high-frequency cortico-muscular coherence (Kristeva-Feige et al., 2002) derives from the rhythmic discharges in the corticospinal neurons projecting to the spinal motoneurons (Farmer et al., 1997) and the increase in functional coupling in infants (Ritterband-Rosenbaum et al., 2017) suggests that the descending beta-band control signal modulates the spinal activity (Lacquaniti et al., 2017), which in turn control muscles through effective low-frequency common drive (Farina, Negro, et al., 2014). Despite the filtering effect performed by muscles (Mannard & Stein, 1973), high-frequency cortical components may also play a role in the force regulation (Watanabe & Kohn, 2015). Co-contraction showed significant EEG-EMG and EMG-EMG coherence in the beta-band among antagonistic muscles, suggesting a direct cortical regulation of both agonist and antagonist muscles (Dal Maso et al., 2017), although fluctuating generation of force would result in a coherence peak in the alpha-band (Hansen et al., 2002; Negro et al., 2009). While the coherence peak identified in the alpha-band in both co-activation conditions identified a common input for force generation, the peak in the beta-band, identified only during co-activation for impedance modulation, confirmed the cortical origin of the co-contraction input.

While voluntary actions are achieved by the recruitment of Ia inhibitory interneurons, which depresses the activity of spinal motoneurons innervating antagonist muscles (J. B. Nielsen, 2004), the co-contraction of antagonist muscles requires a pathway that both facilitates the simultaneous activation of the antagonistic muscles and maintains a low disynaptic reciprocal inhibition of Ia interneurons (Nielsen, J., Kagamihara, 1992). Therefore, our results suggested the existence of two separate descending pathways (J. Nielsen et al., 1993) centrally originated (Y. Nielsen & Kagamihara, 1993), which could be regulated by the cerebellum (Babadi et al., 2021), to drive motoneurons of antagonistic muscle pairs: one activating the agonist while depressing the antagonist, responsible for torque generation, and the other co-activating both muscles, responsible for stiffness modulation (Hansen et al., 2002). These pathways originate in two distinct anatomical portions of the premotor cortex (Haruno et al., 2012), and are regulated by intracortical excitatory connections (Ethier et al., 2007). Moreover, these pathways may project to spinal premotor interneurons innervating motor pools of multiple antagonist muscles (Ronzano et al., 2021) selectively recruited during torque or stiffness generation.

This study suggests the existence of muscle synergies specifically recruited to modulate impedance and encoding co-contraction in their structures. The existence of muscle synergies has been supported by the low-dimensionality observed in the muscle patterns during several tasks (Borzelli et al., 2013; D’Avella et al., 2006; Dominici et al., 2011; Overduin et al., 2012; Torres-Oviedo & Ting, 2010) and by slower learning after EMG-to-force remapping that require new synergies (Borzelli et al., 2013; D’Avella et al., 2006; Dominici et al., 2011; Giszter et al., 2007; Hart & Giszter, 2010; Overduin et al., 2012; Torres-Oviedo & Ting, 2010), by slower learning after EMG-to-force remapping that require new synergies (Berger et al., 2013, 2022; Borzelli et al., 2022), but how a synergistic command drives pools of MNs of different muscles to modulate their firing rates is still unclear. Muscles driven by the same synergy show higher coherence (Danna-Dos-Santos et al., 2014; De Marchis et al., 2015) and pools of MNs of a single (Hug et al., 2021) or two synergistic (Del Vecchio et al., 2022) muscles are driven by different synaptic inputs. A significant beta-band cortico-synergy coherence (Zandvoort et al., 2019) suggests that high-frequency input to the spinal cord encodes the muscle synergy structure. We speculate that the detected high-frequency common input, which occurred only when an increased co-contraction was required, represented those synergies specifically recruited to modulate the joint impedance and encodes co-contraction in their structures.

The residual coherence analysis allowed the identification of two subsets of MNs, roughly 1/3 and 1/4 of those identified respectively on the TB and on the BB, that were selectively recruited either to modulate co-contraction or to generate force through a conservative criterion. While task-specific motor units have been reported before (Borzelli, Gazzoni, et al., 2020; Herrmann & Flanders, 1998; Hodson-Tole et al., 2013; ter Haar Romeny et al., 1984), this study identified, for the first time, a set of units selectively recruited to modulate co-contraction. Taken together, our results suggest the existence of a separate synaptic pathway for impedance modulation, which is cortically originated and projects to a specific pool of motor units.

Our results provide a physiological basis for the exploitation of co-contraction to control the impedance of a robotic device (Ajoudani et al., 2012; Borzelli, Burdet, et al., 2020) and for motor augmentation (Abdi et al., 2016; Dominijanni et al., 2021; Eden et al., 2021; Salvietti et al., 2017). The separate neural pathways driving co-contraction may be exploited as an implicit “task null space” (Lisini Baldi et al., 2021) to control extra degrees of freedom without affecting the control of the natural degree of freedom involved in performing a task. Similarly, the beta-component of the common drive was volitionally modulated to control a cursor in real-time (Bräcklein et al., 2021). The feasibility of the use of co-contraction in motor augmentation was recently demonstrated during an isometric multi-muscle force generation task (Gurgone et al., 2022). Moreover, the proposed approach may be transferred to patients with neuromuscular disorders, such as Stroke (Rosa et al., 2014), Dystonia (Malfait & Sanger, 2007), or Parkinson desease (Fung et al., 2000), to characterize pathological features in the synaptic input to multiple coactive muscles that differs from healthy subjects.

Overall, our study has expanded the current understanding of the control of antagonist muscles for joint impedance modulation demonstrating, for the first time, the existence of separate pathways driving, at different frequency bands, the generation of force or the modulation of impedance by selectively activating different pools of MNs.

## Acknowledgements

This work was supported by the Italian University Ministry (PRIN grant 2015HFWRYY, PRIN grant 2017CBF8NJ, PRIN grant 2020EM9A8X).

## References

Abdi, E., Burdet, E., Bouri, M., Himidan, S., & Bleuler, H. (2016). In a demanding task, three-handed manipulation is preferred to two-handed manipulation. Scientific Reports, 6. https://doi.org/10.1038/srep21758

Ajoudani, A., Tsagarakis, N. G., & Bicchi, A. (2012). Tele-impedance: Towards transferring human impedance regulation skills to robots. 2012 IEEE International Conference on Robotics and Automation, 382–388. https://doi.org/10.1109/ICRA.2012.6224904

Babadi, S., Vahdat, S., & Milner, T. E. (2021). Neural Substrates of Muscle Co-contraction during Dynamic Motor Adaptation. Journal of Neuroscience, 41(26), 5667–5676. https://doi.org/10.1523/JNEUROSCI.2924-19.2021

Berger, D. J., Borzelli, D., & d’Avella, A. (2022). Task space exploration improves adaptation after incompatible virtual surgeries. Journal of Neurophysiology, 127(4), 1127–1146. https://doi.org/10.1152/jn.00356.2021

Berger, D. J., Gentner, R., Edmunds, T., Pai, D. K., & d’Avella, A. (2013). Differences in Adaptation Rates after Virtual Surgeries Provide Direct Evidence for Modularity. Journal of Neuroscience, 33(30), 12384–12394. https://doi.org/10.1523/JNEUROSCI.0122-13.2013

Bernstein, N. (1967). The co-ordination and regulation of movements.

Berret, B., & Jean, F. (2020). Stochastic optimal open-loop control as a theory of force and impedance planning via muscle co-contraction. PLoS Computational Biology. https://doi.org/10.1371/journal.pcbi.1007414

Borzelli, D., Berger, D. J., Pai, D. K., & D’Avella, A. (2013). Effort minimization and synergistic muscle recruitment for three-dimensional force generation. Frontiers in computational neuroscience, 7, 186.

Borzelli, D., Burdet, E., Pastorelli, S., d’ Avella, A., & Gastaldi, L. (2020). Identification of the best strategy to command variable stiffness using electromyographic signals. 17(1), 016058. https://doi.org/10.1088/1741-2552/ab6d88

Borzelli, D., Cesqui, B., Berger, D. J., Burdet, E., & D’Avella, A. (2018). Muscle patterns underlying voluntary modulation of co-contraction. PLOS ONE, 13(10), e0205911. https://doi.org/10.1371/journal.pone.0205911

Borzelli, D., Gazzoni, M., Botter, A., Gastaldi, L., D’Avella, A., & Vieira, T. M. (2020). Contraction level, but not force direction or wrist position, affects the spatial distribution of motor unit recruitment in the biceps brachii muscle. European Journal of Applied Physiology, 120(4), 853–860. https://doi.org/10.1007/s00421-020-04324-6

Borzelli, D., Gurgone, S., Mezzetti, M., De Pasquale, P., Berger, D. J., Milardi, D., Acri, G., & D’Avella, A. (2022). Adaptation to Virtual Surgeries Across Multiple Practice Sessions. In D. Torricelli, M. Akay, & J. L. Pons (A c. Di), Converging Clinical and Engineering Research on Neurorehabilitation IV (pagg. 563–568). Springer International Publishing. https://doi.org/10.1007/978-3-030-70316-5_90

Bräcklein, M., Ibáñez, J., Barsakcioglu, D. Y., & Farina, D. (2021). Towards human motor augmentation by voluntary decoupling beta activity in the neural drive to muscle and force production. Journal of Neural Engineering, 18(1), 016001. https://doi.org/10.1088/1741-2552/abcdbf

Burdet, E., Osu, R., Franklin, D., Milner, T., & Kawato, M. (2001). The central nervous system stabilizes unstable dynamics by learning optimal impedance. Nature, 414, 446–449.

Carter, G. C. (1987). Coherence and time delay estimation. Proceedings of the IEEE, 75(2), 236–255. https://doi.org/10.1109/PROC.1987.13723

Dal Maso, F., Longcamp, M., Cremoux, S., & Amarantini, D. (2017). Effect of training status on beta-range corticomuscular coherence in agonist vs. Antagonist muscles during isometric knee contractions. Experimental Brain Research, 235(10), 3023–3031. https://doi.org/10.1007/s00221-017-5035-z

Danna-Dos-Santos, A., Boonstra, T. W., Degani, A. M., Cardoso, V. S., Magalhaes, A. T., Mochizuki, L., & Leonard, C. T. (2014). Multi-muscle control during bipedal stance: An EMG–EMG analysis approach. Experimental Brain Research, 232(1), 75–87. https://doi.org/10.1007/s00221-013-3721-z

Datta, B. Y. A. K., & Stephens, J. A. (1990). Synchronization of Motor Unit Activity During Volntary Contraction in Man. 397–419.

D’Avella, A., Portone, A., Fernandez, L., & Lacquaniti, F. (2006). Control of fast-reaching movements by muscle synergy combinations. Journal of Neuroscience, 26(30), 7791–7810. https://doi.org/10.1523/JNEUROSCI.0830-06.2006

De Luca, C. J., & Erim, Z. (1994). Common drive of motor units in regulation of muscle force. In Trends in Neurosciences (Vol. 17, Numero 7, pagg. 299–305). https://doi.org/10.1016/0166-2236(94)90064-7

De Luca, C. J., & Mambrito, B. (1987). Voluntary control of motor units in human antagonist muscles: Coactivation and reciprocal activation. Journal of Neurophysiology, 58(3), 525–542. https://doi.org/10.1152/jn.1987.58.3.525

De Marchis, C., Severini, G., Castronovo, A. M., Schmid, M., & Conforto, S. (2015). Intermuscular coherence contributions in synergistic muscles during pedaling. Experimental Brain Research, 233(6), 1907–1919. https://doi.org/10.1007/s00221-015-4262-4

De Serres, S. J., & Milner, T. E. (1991). Wrist muscle activation patterns and stiffness associated with stable and unstable mechanical loads. Experimental Brain Research, 86(2), 451–458. https://doi.org/10.1007/BF00228972

Del Vecchio, A., Germer, C., Kinfe, T. M., Nuccio, S., Hug, F., Eskofier, B., Farina, D., & Enoka, R. M. (2022). Common synaptic inputs are not distributed homogeneously among the motor neurons that innervate synergistic muscles [Preprint]. Neuroscience. https://doi.org/10.1101/2022.01.23.477379

Del Vecchio, A., Germer, C. M., Elias, L. A., Fu, Q., Fine, J., Santello, M., & Farina, D. (2019). The human central nervous system transmits common synaptic inputs to distinct motor neuron pools during non-synergistic digit actions. Journal of Physiology, 597(24), 5935–5948. https://doi.org/10.1113/JP278623

Del Vecchio, A., Holobar, A., Falla, D., Felici, F., Enoka, R. M., & Farina, D. (2020). Tutorial: Analysis of motor unit discharge characteristics from high-density surface EMG signals. Journal of Electromyography and Kinesiology, 53, 102426. https://doi.org/10.1016/J.JELEKIN.2020.102426

Dideriksen, J. L., Negro, F., Falla, D., Kristensen, S. R., Mrachacz-Kersting, N., & Farina, D. (2018). Coherence of the Surface EMG and Common Synaptic Input to Motor Neurons. Frontiers in Human Neuroscience, 12. https://www.frontiersin.org/articles/10.3389/fnhum.2018.00207

Dominici, N., Ivanenko, Y. P., Cappellini, G., d’Avella, A., Mondì, V., Cicchese, M., Fabiano, A., Silei, T., Di Paolo, A., Giannini, C., Poppele, R. E., & Lacquaniti, F. (2011). Locomotor primitives in newborn babies and their development. Science (New York, N.Y.), 334(6058), 997–999. https://doi.org/10.1126/science.1210617

Dominijanni, G., Shokur, S., Salvietti, G., Buehler, S., Palmerini, E., Rossi, S., De Vignemont, F., d’Avella, A., Makin, T. R., Prattichizzo, D., & Micera, S. (2021). The neural resource allocation problem when enhancing human bodies with extra robotic limbs. In Nature Machine Intelligence. https://doi.org/10.1038/s42256-021-00398-9

Eden, J., Bracklein, M., Ibanez Pereda, Jaime Barsakcioglu, Deren Yusuf Di Pino, G., Farina, G., Burdet, E., & Mehring, C. (2021). Human movement augmentation and how to make it a reality. arXiv, 2106.08129.

Enoka, R. M., & Fuglevand, A. J. (2001). Motor unit physiology: Some unresolved issues. Muscle & Nerve, 24(1), 4–17. https://doi.org/10.1002/1097-4598(200101)24:1<4::AID-MUS13>3.0.CO;2-F

Ethier, C., Brizzi, L., Gigue, D., Sperimentale, M., & Umana, F. (2007). Corticospinal control of antagonistic muscles in the cat. European Journal of Neuroscience, 26(6), 1632–1641. https://doi.org/10.1111/j.1460-9568.2007.05778.x

Farina, D., Merletti, R., & Enoka, R. M. (2014). The extraction of neural strategies from the surface EMG: An update. Journal of Applied Physiology, 117(11), 1215–1230. https://doi.org/10.1152/japplphysiol.00162.2014

Farina, D., & Negro, F. (2015). Common synaptic input to motor neurons, motor unit synchronization, and force control. Exercise and Sport Sciences Reviews, 43(1), 23–33. https://doi.org/10.1249/JES.0000000000000032

Farina, D., Negro, F., & Dideriksen, J. L. (2014). The effective neural drive to muscles is the common synaptic input to motor neurons. Journal of Physiology, 592(16), 3427–3441. https://doi.org/10.1113/jphysiol.2014.273581

Farmer, S. F., Bremner, F. D., Halliday, D. M., Rosenberg, J. R., & Stephens, J. A. (1993). The frequency content of common synaptic inputs to motoneurones studied during voluntary isometric contraction in man. The Journal of Physiology, 470(1), 127–155. https://doi.org/10.1113/jphysiol.1993.sp019851

Farmer, S. F., Halliday, D. M., Conway, B. A., Stephens, J. A., & Rosenberg, J. R. (1997). A review of recent applications of cross-correlation methodologies to human motor unit recording. Journal of Neuroscience Methods, 74(2), 175–187. https://doi.org/10.1016/S0165-0270(97)02248-6

Forster, E., Simon, U., Augat, P., & Claes, L. (2004). Extension of a state-of-the-art optimization criterion to predict co-contraction. Journal of Biomechanics. https://doi.org/10.1016/j.jbiomech.2003.09.003

Freund, H. J. (1983). Motor unit and muscle activity in voluntary motor control. https://doi.org/10.1152/physrev.1983.63.2.387, 63(2), 387–436. https://doi.org/10.1152/PHYSREV.1983.63.2.387

Fung, V. S. C., Burne, J. A., & Morris, J. G. L. (2000). Objective quantification of resting and activated parkinsonian rigidity: A comparison of angular impulse and work scores. Movement Disorders, 15(1), 48–55. https://doi.org/10.1002/1531-8257(200001)15:1<48::AID-MDS1009>3.0.∞;2-E

Giszter, S., Patil, V., & Hart, C. (2007). Primitives, premotor drives, and pattern generation: A combined computational and neuroethological perspective. Progress in Brain Research, 165, 323–346. Scopus. https://doi.org/10.1016/S0079-6123(06)65020-6

Gribble, P. L., Mullin, L. I., Cothros, N., & Mattar, A. (2003). Role of cocontraction in arm movement accuracy. Journal of Neurophysiology, 89(5), 2396–2405.

Gurgone, S., Borzelli, D., Pasquale, P. de, Berger, D. J., Baldi, T. L., D’Aurizio, N., Prattichizzo, D., & d’Avella, A. (2022). Simultaneous control of natural and extra degrees of freedom by isometric force and electromyographic activity in the muscle-to-force null space. Journal of Neural Engineering, 19(1), 016004. https://doi.org/10.1088/1741-2552/ac47db

Hansen, S., Hansen, N. L., Christensen, · L O D, Petersen, · N T, & Nielsen, J. B. (2002). Coupling of antagonistic ankle muscles during co-contraction in humans. Exp Brain Res, 146(3), 282–292. https://doi.org/10.1007/s00221-002-1152-3

Hart, C. B., & Giszter, S. F. (2010). A Neural Basis for Motor Primitives in the Spinal Cord. Journal of Neuroscience, 30(4), 1322–1336. https://doi.org/10.1523/JNEUROSCI.5894-08.2010

Haruno, M., Ganesh, G., Burdet, E., & Kawato, M. (2012). Differential neural correlates of reciprocal activation and cocontraction control in dorsal and ventral premotor cortices. Journal of Neurophysiology, 107(1), 126–133. https://doi.org/10.1152/JN.00735.2010/ASSET/IMAGES/LARGE/Z9K0011211200005.JPEG

Haynes, E. M. K., & Kim, C. (2021). Antagonist surface electromyogram decomposition and the case of the missing motor units. Journal of Neurophysiology, 126(6), 1943–1947. https://doi.org/10.1152/jn.00435.2021

Hermens, H. J., Freriks, B., Merletti, R., Stegeman, D., Blok, J., Rau, G., Disselhorst-Klug, C., & Hägg, G. (1999). European Recommendations for Surface ElectroMyoGraphy Results of the SENIAM project. In Roessingh Research and Development.

Herrmann, U., & Flanders, M. (1998). Directional Tuning of Single Motor Units. J. Neurosci., 18(20), 8402–8416.

Hodson-Tole, E. F., Loram, I. D., & Vieira, T. M. M. (2013). Myoelectric activity along human gastrocnemius medialis: Different spatial distributions of postural and electrically elicited surface potentials. Journal of Electromyography and Kinesiology, 23(1), 43–50. https://doi.org/10.1016/j.jelekin.2012.08.003

Hogan, N. (1984). Adaptive Control of Mechanical Impedance by Coactivation of Antagonist Muscles. IEEE Transactions on Automatic Control. https://doi.org/10.1109/TAC.1984.1103644

Holobar, A., & Zazula, D. (2007). Multichannel blind source separation using convolution Kernel compensation. IEEE Transactions on Signal Processing. https://doi.org/10.1109/TSP.2007.896108

Hug, F., Del Vecchio, A., Avrillon, S., Farina, D., & Tucker, K. (2021). Muscles from the same muscle group do not necessarily share common drive: Evidence from the human triceps surae. Journal of Applied Physiology. https://doi.org/10.1152/JAPPLPHYSIOL.00635.2020

Hughes, R. E., Bean, J. C., & Chaffin, D. B. (1995). Evaluating the effect of co-contraction in optimization models. Journal of Biomechanics. https://doi.org/10.1016/0021-9290(95)95277-C

Kendall, F., McCreary, E., & Provance, P. (1994). Muscles, Testing and Function. Medicine and Science in Sports and Exercise, 26(8), 1070. https://doi.org/10.1249/00005768-199408000-00023

Kristeva-Feige, R., Fritsch, C., Timmer, J., & Lücking, C. H. (2002). Effects of attention and precision of exerted force on beta range EEG-EMG synchronization during a maintained motor contraction task. Clinical Neurophysiology, 113(1), 124–131. https://doi.org/10.1016/S1388-2457(01)00722-2

Lacquaniti, F., & Maioli, C. (1989). The role of preparation in tuning anticipatory and reflex responses during catching. Journal of Neuroscience, 9(1), 134.

Lacquaniti, F., Zago, M., & Farina, D. (2017). Tick-tock, spinal motor neurons go with the cortical clock in young infants. J Physiol, 595, 2405–2406. https://doi.org/10.1113/JP273901

Laine, C. M., Martinez-Valdes, E., Falla, D., Mayer, F., & Farina, D. (2015). Motor neuron pools of synergistic thigh muscles share most of their synaptic input. Journal of Neuroscience, 35(35), 12207–12216. https://doi.org/10.1523/JNEUROSCI.0240-15.2015

Latash, M. (1992). Independent control of joint stiffness in the framework of the equilibrium-point hypothesis. Biological cybernetics, 67(4), 377–384.

Lisini Baldi, T., D’Aurizio, N., Gaudeni, C., Gurgone, S., Borzelli, D., D’Avella, A., & Prattichizzo, D. (2021). Exploiting Implicit Kinematic Kernel forControlling a Wearable Robotic Extra-finger. arXiv, 2012.03600.

Malfait, N., & Sanger, T. D. (2007). Does dystonia always include co-contraction? A study of unconstrained reaching in children with primary and secondary dystonia. Experimental Brain Research, 176(2), 206–216. https://doi.org/10.1007/s00221-006-0606-4

Mannard, A., & Stein, R. B. (1973). Determination of the frequency response of isometric soleus muscle in the cat using random nerve stimulation. The Journal of Physiology, 229(2), 275–296. https://doi.org/10.1113/jphysiol.1973.sp010138

Martinez-Valdes, E., Negro, F., Falla, D., De Nunzio, A. M., & Farina, D. (2018). Surface electromyographic amplitude does not identify differences in neural drive to synergistic muscles. Journal of Applied Physiology, 124(4), 1071–1079. https://doi.org/10.1152/japplphysiol.01115.2017

Milner, T. (2002). Adaptation to destabilizing dynamics by means of muscle cocontraction. Experimental Brain Research.

Nandi, T., Hortobágyi, T., van Keeken, H. G., Salem, G. J., & Lamoth, C. J. C. (2019). Standing task difficulty related increase in agonist-agonist and agonist-antagonist common inputs are driven by corticospinal and subcortical inputs respectively. Scientific Reports, 9(1), 2439. https://doi.org/10.1038/s41598-019-39197-z

Negro, F., Holobar, A., & Farina, D. (2009). Fluctuations in isometric muscle force can be described by one linear projection of low-frequency components of motor unit discharge rates. J Physiol, 14.

Nielsen, J. B. (2004). Sensorimotor integration at spinal level as a basis for muscle coordination during voluntary movement in humans. J Appl Physiol, 96(5), 1961–1967. https://doi.org/10.1152/japplphysiol.01073.2003.-Spinal

Nielsen, J., Kagamihara, Y. (1992). The regulation of disynaptic reciprocal Ia inhibition during co-contraction of antagonistic muscles in man. The Journal of physiology, 456(1), 373–391.

Nielsen, J., Petersen, N., Deuschlt, G., & Ballegaard, M. (1993). TASK-RELATED CHANGES IN THE EFFECT OF MAGNETIC BRAIN STIMULATION ON SPINAL NEURONES IN MAN. In Journal of Physiology (Vol. 471).

Nielsen, Y., & Kagamihara, J. (1993). The regulation of presynaptic inhibition during co-contraction of antagonistic muscles in man. The Journal of physiology, 464(1), 575–593.

Ohtsuka, H., Nakajima, T., Komiyama, T., Suzuki, S., Irie, S., & Ariyasu, R. (2022). Execution of natural manipulation in the air enhances the beta-rhythm intermuscular coherences of the human arm depending on muscle pairs. Journal of Neurophysiology, 127(4), 946–957. https://doi.org/10.1152/jn.00421.2021

Overduin, S. A., d’Avella, A., Carmena, J. M., & Bizzi, E. (2012). Microstimulation activates a handful of muscle synergies. Neuron, 76(6), 1071–1077. https://doi.org/10.1016/j.neuron.2012.10.018

Power, K. E., Lockyer, E. J., Botter, A., Vieira, T., & Button, D. C. (2022). Endurance-exercise training adaptations in spinal motoneurones: Potential functional relevance to locomotor output and assessment in humans. European Journal of Applied Physiology, 122(6), 1367–1381. https://doi.org/10.1007/s00421-022-04918-2

Ritterband-Rosenbaum, A., Herskind, A., Li, X., Willerslev-Olsen, M., Damgaard Olsen, M., Farmer, S. F., Nielsen, J. B., & Nielsen, J. B. (2017). A critical period of corticomuscular and EMG-EMG coherence detection in healthy infants aged 9-25 weeks. The Journal of Physiology C 2016 The Authors. The Journal of Physiology C, 595(8), 2699–2713. https://doi.org/10.1113/JP273090

Ronzano, R., Lancelin, C., Bhumbra, G. S., Brownstone, R. M., & Beato, M. (2021). Proximal and distal spinal neurons innervating multiple synergist and antagonist motor pools. eLife, 10. https://doi.org/10.7554/ELIFE.70858

Rosa, M. C. N., Marques, A., Demain, S., & Metcalf, C. D. (2014). Lower limb co-contraction during walking in subjects with stroke: A systematic review. Journal of Electromyography and Kinesiology, 24(1), 1–10. https://doi.org/10.1016/j.jelekin.2013.10.016

Rosenberg, J. R., Amjad, A. M., Breeze, P., Brillinger, D. R., & Halliday, D. M. (1989). The Fourier approach to the identification of functional coupling between neuronal spike trains. In Progress in Biophysics and Molecular Biology (Vol. 53, Numero 1, pagg. 1–31). https://doi.org/10.1016/0079-6107(89)90004-7

Rosenberg, J. R., Halliday, D. M., Breeze, P., & Conway, B. A. (1998). Identification of patterns of neuronal connectivity—Partial spectra, partial coherence, and neuronal interactions. Journal of Neuroscience Methods, 83(1), 57–72. https://doi.org/10.1016/S0165-0270(98)00061-2

Salvietti, G., Hussain, I., Cioncoloni, D., Taddei, S., Rossi, S., & Prattichizzo, D. (2017). Compensating Hand Function in Chronic Stroke Patients Through the Robotic Sixth Finger. IEEE Transactions on Neural Systems and Rehabilitation Engineering, 25(2), 142–150. https://doi.org/10.1109/TNSRE.2016.2529684

Selen, L. P. J., Beek, P. J., & Van Dieën, J. H. (2006). Impedance is modulated to meet accuracy demands during goal-directed arm movements. Experimental Brain Research, 172(1), 129–138. https://doi.org/10.1007/s00221-005-0320-7

Selen, L. P. J., Franklin, D. W., & Wolpert, D. M. (2009). Impedance control reduces instability that arises from motor noise. Journal of Neuroscience, 29(40), 12606–12616.

Takagi, A., Kambara, H., & Koike, Y. (2020). Independent control of cocontraction and reciprocal activity during goal-directed reaching in muscle space. Scientific Reports. https://doi.org/10.1038/s41598-020-79526-1

ter Haar Romeny, B. M., van der Gon, J. J., & Gielen, C. C. (1984). Relation between location of a motor unit in the human biceps brachii and its critical firing levels for different tasks. Experimental neurology, 85(3), 631–650.

Torres-Oviedo, G., & Ting, L. H. (2010). Subject-Specific Muscle Synergies in Human Balance Control Are Consistent Across Different Biomechanical Contexts. Journal of Neurophysiology, 103(6), 3084–3098. https://doi.org/10.1152/jn.00960.2009

Vieira, T. M., & Botter, A. (2021). The Accurate Assessment of Muscle Excitation Requires the Detection of Multiple Surface Electromyograms. Exercise and Sport Sciences Reviews. https://doi.org/10.1249/JES.0000000000000240

Watanabe, R. N., & Kohn, A. F. (2015). Fast Oscillatory Commands from the Motor Cortex Can Be Decoded by the Spinal Cord for Force Control. The Journal of Neuroscience, 35(40), 13687–13697. https://doi.org/10.1523/JNEUROSCI.1950-15.2015

Zandvoort, C. S., van Dieën, J. H., Dominici, N., & Daffertshofer, A. (2019). The human sensorimotor cortex fosters muscle synergies through cortico-synergy coherence. NeuroImage, 199, 30–37. https://doi.org/10.1016/j.neuroimage.2019.05.041

